# Two δ-Catenins, Plakophilin 4 and p120, Promote Formation of Distinct Types of Adherens Junctions

**DOI:** 10.1101/2025.06.14.659701

**Authors:** Indrajyoti Indra, Regina B. Troyanovsky, Farida V. Korobova, Sergey M. Troyanovsky

**Affiliations:** Department of Dermatology, Northwestern University, The Feinberg School of Medicine, Chicago, IL 60611; Center for Advanced Microscopy/Nikon Imaging Center, Northwestern University, Chicago, IL 60614; Department of Cell & Developmental Biology, Northwestern University, The Feinberg School of Medicine, Chicago, IL 60614

**Keywords:** <u>Key words</u>: cadherin, p120-catenin, plakophilin 4, binding to actin, adherens junctions

## Abstract

Classic cadherins are instrumental for joining cells into tissues by producing cell-cell adhesions known as adherens junctions (AJs). These morphologically diverse structures are tailored to the specific cell sites, type of cells, and particular functions. The mechanism of AJ diversification remains unknown. Here we show that two members of the δ-catenin protein family, p120 and plakophilin 4 (pkp4), which interact with the juxtamembrane intracellular region of classic cadherins, promote distinct types of cadherin clustering thereby contributing to AJ specialization. The type controlled by p120 is driven by interactions between cadherin-associated protein, α-catenin, and actin filaments. This “canonical” clustering mechanism results in formation of AJs that play a major role in overall cell-cell adhesion. The type promoted by pkp4 is driven by an α-catenin-independent cadherin-F-actin interaction. It generates the so-called lateral spot AJs, which apparently function in processes other than cell-cell adhesion. Collectively, our study shows how δ-catenins regulate a balance between different types of AJs in epithelial cells.

## Introduction

Adherens Junctions (AJs) are cell-cell adhesive structures interconnecting cells in nearly all multicellular organisms [1–4]. In addition to the prime role in adhesion, AJs are also involved in myriad signal transduction pathways including force sensing [5–8]. The mechanisms that support a remarkable functional diversity of AJs remain to be understood.

AJs are also incredibly variable in terms of morphology, localization, and repertoire of the accessory proteins [9–11]. The most well studied AJs, apical AJs, are formed at the apical region of the lateral plasma membrane. They, in turn, could be subdivided into the linear AJs (also known as *zonula adhaerens*) and the punctate AJs (known also as *fascia adhaerens*, focal, or radial AJs). A second type of AJs, basal AJs, are morphologically similar to the punctate apical AJs, but form at the basal portion of the lateral membrane [12, 13]. The basal AJs, in contrast to the relatively immobile apical AJs, exhibit upward directional movement [14, 15].

The apical linear AJs (apical-lAJs) and apical or basal punctate AJs (apical/basal-pAJs) play a major role in intercellular adhesion by interconnecting the actomyosin cytoskeleton of individual cells within the tissues. In addition to these widely studied types of AJs, a middle section of the cell-cell contacts in most epithelia exhibits a third, largely neglected type of AJs, “spot-like” AJs (lateral sAJs) of a submicron size. While these AJs (also known as lateral AJs or *puncta adhaerentia*) also interact with actin filaments, they neither associate with vinculin and afadin, the two key actin-binding proteins of the apical and basal AJs, nor they show any directional movement [12, 14, 16, 17]. The specific enrichment of these AJs with signaling proteins, such as erbin [18] or PLEKHA5 [19] together with some functional experiments [17] suggest that lateral-sAJs could be implicated in the intercellular signaling events. General mechanisms underlying AJ specialization have not been studied.

A key structural unit of all types of AJs is the cadherin-catenin complex (CCC). Its transmembrane receptor (E-cadherin in epithelia) belongs to the family of classic cadherins. The intracellular tail of these cadherins associates with two Armadillo (ARM)-repeat proteins [1–4]. A distal, C-terminal region of the tail interacts with a 12 ARM repeat protein β-catenin (or its ortholog, plakoglobin), which links cadherins to the actin-binding protein α-catenin. The juxtamembrane region of the tail interacts with p120-catenin (p120) or its orthologs, plakophilin 4 (pkp4), δ2-catenin, and ARVCF, which comprise a separate subfamily, δ-catenins, of 9 ARM repeat proteins. CCC-associated proteins could play a leading role in directing cadherin into functionally and morphologically distinct types of AJs [20].

Here we tested the roles of p120 and pkp4 in AJ diversification. These two proteins evolved from a common ancestor at the beginning of vertebrate evolution [21, 22]. Majority of epithelial cells in mammals co-express these two proteins together with their relative, ARVCF. Since these ARM repeat proteins bind to the same site of E-cadherin [23], they form three different types of CCC, denoted below as p120-CCC, pkp4-CCC, and ARVCF-CCC. In most epithelial cells the p120-CCC appeared to be a predominant form. The δ-catenin proteins have been known to complement one another in two important functions: they regulate the lifetime of the CCC by controlling the accessibility of cadherin endocytic motifs [24, 25] and modulate the strength and dynamics of AJs by controlling the AJ-associated cytoskeleton through Rho GTPase signaling [26, 27]. It is assumed that distinct δ-catenins may specifically tune these regulatory mechanisms [28, 29]. However, an immunomorphological observation from Dr. Franke’s laboratory showing that pkp4 is enriched only in a subset of AJs [30] has provided a hint that the proteins of this family might be involved in AJ diversification. The immunolocalization and functional analyses of p120 and pkp4 reported here present direct evidence in support of this assumption. Furthermore, our data suggest that these proteins define a balance between distinct morphological types of AJs by promoting two specialized pathways of AJ assembly. While p120-CCC facilitates formation of the apical/basal-pAJs, where CCC clustering is based on the well-studied α-catenin-actin interactions, pkp4-CCC promotes formation of the lateral-sAJs, where cadherin clustering is reinforced by a novel, α-catenin independent interaction with actin filaments. Thus, a competition between canonical, p120-maintained, and noncanonical, pkp4-maintained, pathways of AJ assembly contributes to the spatial organization of cell-cell adhesions.

## Results

### Pkp4 is predominantly recruited into lateral-sAJs

Epidermal A431 cells, as most other human epithelial cells, simultaneously express three δ-catenins thus forming p120-CCC, pkp4-CCC, and ARVCF-CCC. Proteomics analysis of A431 cells suggested that the latter two species account for only about 20% of all CCCs [20]. Here we focused on p120-CCC and pkp4-CCC. Triple staining of A431 cells for pkp4, p120, and E-cadherin showed that pkp4-CCC was not fully colocalized with p120-CCC but was enriched in distinct AJs or in separate clusters within the larger AJs (Fig. 1a, b). Furthermore, detail inspection of the basal, middle, and apical confocal sections revealed a remarkable feature: pkp4 appeared to be most prominent in AJs located at the middle portion of the lateral membrane, which corresponded to the lateral-sAJs by their morphology (Fig. 1c). By contrast, p120, while also observed in the lateral-sAJs, was especially enriched in pAJs at both apical and basal locations (apical-and basal-pAJs). Such spatial segregation of two CCCs was further underscored by scatterplot analysis of fluorophore pixel intensities of the obtained images (Fig. 1b, right panel). It showed much higher linear relationship between p120 and E-cadherin (Pearson’s correlation coefficient, r ∼ 0.85), than that between pkp4 and E-cadherin (r ∼ 0.55) or pkp4 and p120 (r ∼ 0.45). In the latter case only a small fraction of Pkp4-and p120-derived pixels in the scatterplots were interdependent (Fig. 1b).

**Figure 1.**
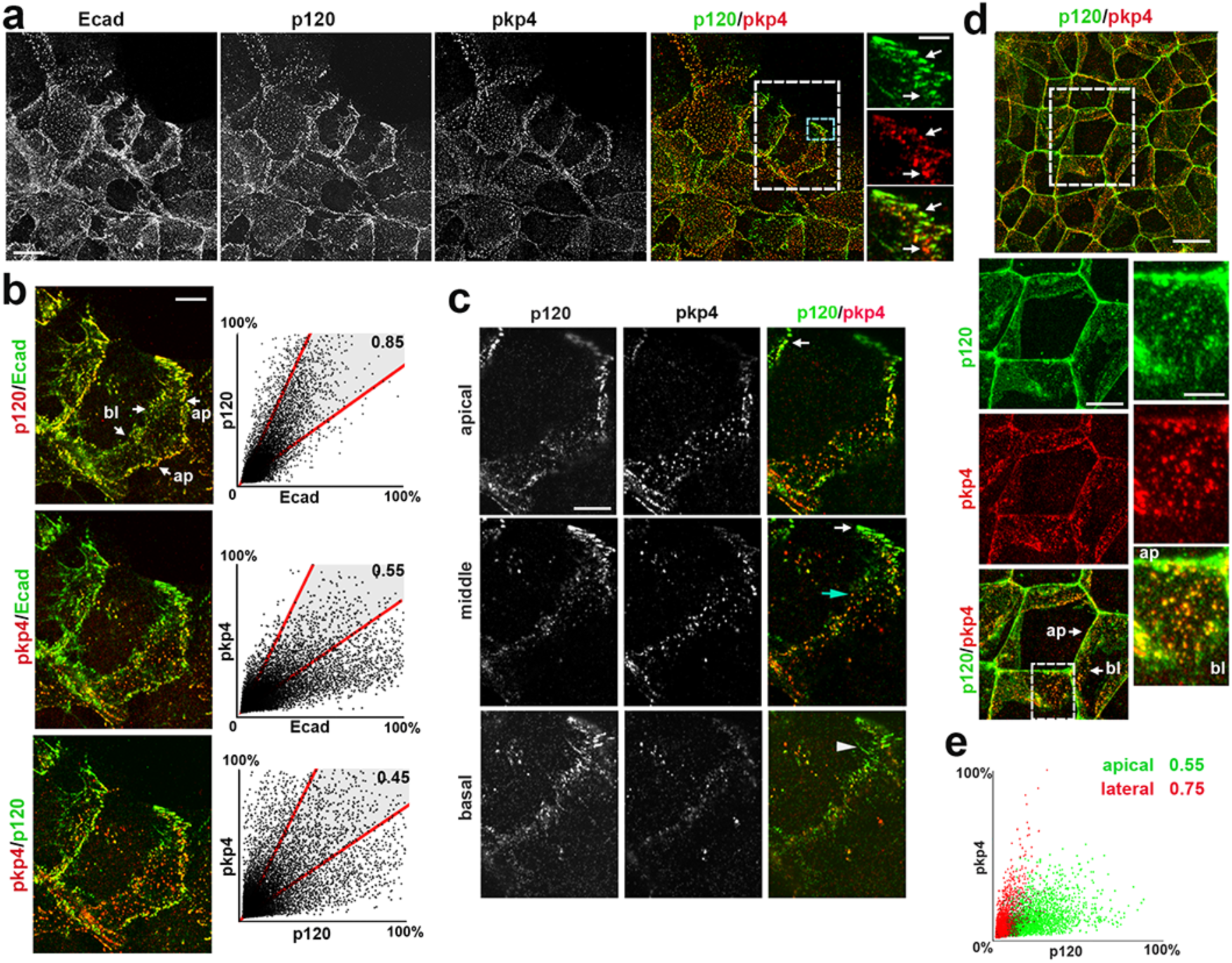
Lateral and apical AJs in A431 and DLD cells recruit different combinations of p120 and pkp4. (**a**) Projections of all x-y optical slices of A431 cells triple stained for E-cadherin (Ecad), p120 (p120), and pkp4 (pkp4). The merged image shows only p120 (green) and pkp4 (red) staining. Bar, 20 μm. A blue dashed boxed area is enlarged at the right. The arrows point at the pkp4-enriched clusters, which could completely lack p120 (the bottom arrow) or be embedded into the p120-rich pAJs (the upper arrow). Bar, 5 µm. (**b**) The zoomed area marked by the dashed white box in (a) is presented in three different staining combinations: p120 (red)/Ecad (green), pkp4 (red)/Ecad (green) and pkp4 (red)/p120 (green). Note that both p120 and E-cadherin are present in nearly all AJs while pkp4 mostly resides in the dot-like AJs. Bar, 20 μm. Apical (ap) and basal (bl) ends of the lateral membranes are indicated. The corresponding scatterplots of red and green relative pixel intensities are at the right panel. The r values are shown at the upper right. Note that p120/Ecad pixels are clustered within a zone of high positive relationship (indicated by shaded area between the red lines). The pkp4/Ecad or pkp4/p120 pixels show lower correlations. (**c**) Projections of optical z slices spanning apical, middle, and basal portions of the central cell shown in (b). Bar, 10 μm. The apical region is reconstructed by a projection spanning the apical 0.9 µm of cells, the middle region is a projection spanning next 1.5 µm, and the basal region is a projection spanning 0.6 µm of the cell base. Bar, 20 μm. Note that p120 is mostly present in the apical AJs (white arrows) and basal AJs (arrowhead), while pkp4 is enriched in dot-like AJs at the middle cell-cell contact area (one of which is indicated by a blue arrow). (**d**) Projections of all x-y optical slices of DLD1 cells stained for p120 (green), and pkp4 (red). Only merged image is shown. Bar, 20 μm. The zoomed view of the dashed boxed area is presented at the bottom left. Bar, 10 μm. Its dashed boxed area is further zoomed at the right. Bar, 5 μm. (**e**) Scatterplots of p120/pkp4 relative pixel intensities of the apical (green) and lateral (red) cell slices of the cell shown in (d). The r values for each population of pixels are indicated at the upper left.

To verify that a preference of pkp4-CCC for the lateral-sAJs is a general phenomenon for epithelial cells, we investigated colon carcinoma cells, DLD1. Double staining of these cells for pkp4 and p120 also showed that pkp4 is abundant in the lateral-sAJs, while p120 is a dominant species in the apical-lAJs (Fig. 1d), which associates in these cells with the circular actin bundle [18]. Accordingly, scatterplots of pkp4 and p120 pixel intensities collected from the apical and lateral focal planes (Fig. 1e) showed two different linear relationships (with r∼055 and ∼0.75, correspondingly).

### Lateral-sAJs and apical/basal-pAJs incorporate distinct sets of proteins

Previous characterization of pAJs in A431 cells have shown that these AJs are enriched with the tension-dependent actin-binding proteins such as vinculin and afadin [12, 31]. Thus, to validate that p120 and pkp4 favored distinct types of AJs, we studied their co-localization with these tension-dependent proteins. Visual inspection of the stained cells confirmed that vinculin and afadin were mostly associated with the p120-enriched AJs (Fig. 2a). In agreement with this, the scatterplots of red (vinculin or afadin) and green (p120 or pkp4) pixel intensities clearly showed that a significant fraction of pixels derived from the force-sensitive proteins correlated with p120-derived pixels. By contrast, such correlation was not observed between these proteins and pkp4 (Fig. 2c). Accordingly, this group of proteins showed relatively low Pearson’s correlation value with pkp4 (r< 0.5), while it was above 0.5 in case of p120 (Fig. 2c,e).

**Figure 2.**
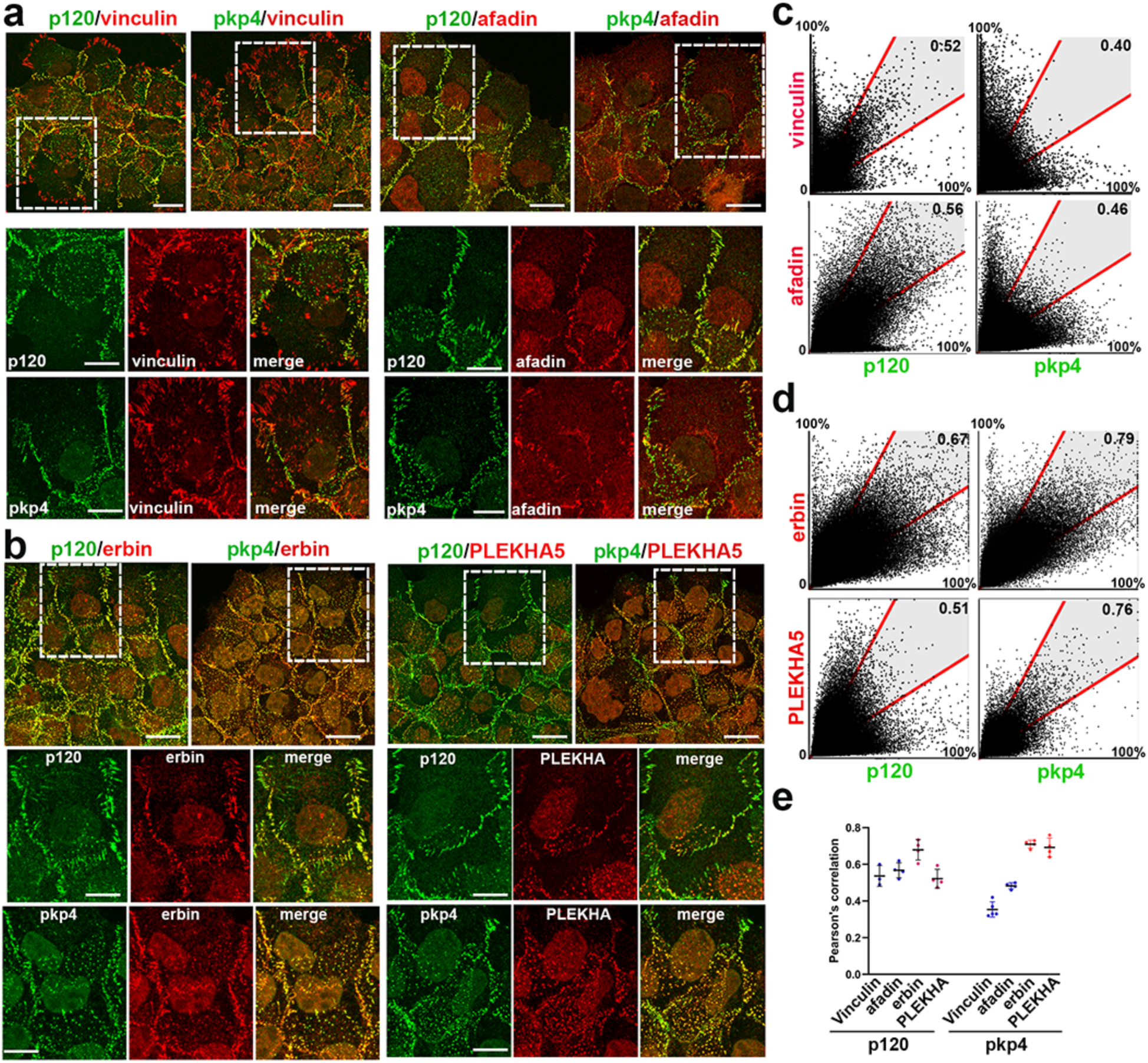
Pkp4-and p120-enriched AJs associate with different sets of proteins. (**a, b**) Projections of all x-y optical slices of A431 cells double stained for p120 or pkp4 (green) in combination with apical/basal-pAJ markers, vinculin and afadin (**a**) or lateral-sAJ markers, erbin, and PLEKHA5 (**b**). Only merged images are shown for low magnifications. Bar, 20 μm. The zoomed areas marked by the white dashed box in the merged images are shown in both colors at the bottom. Bar, 12 μm. (**c, d**) The corresponding scatterplots of red and green relative pixel intensities of the images shown in **a** and **b**. The corresponding r values are shown at the upper right. Note that significant populations of p120/vinculin and p120/afadin pixels show high positive relationship indicated by shaded area between the red lines. Such pools of positively related pixels are undetectable in case of pkp4/vinculin or pkp4/afadin combination (**c**). By contrast, pkp4 shows better relationship with the sAJs markers, PLEKHA5 and erbin (**d**). (**e**) Average Pearson’s colocalization values (r) of images stained as in **a** and **b**. Four independent images for each combination were quantified. Note that combinations between pAJ markers (vinculin and afadin) with p120 show much higher r values than with pkp4, and the relationship is opposite in case of the lateral-sAJ markers (PLEKHA5 and erbin).

While specific protein composition of the lateral-sAJs had not been accurately investigated, available data suggest that they are enriched with erbin [18] and PLEKHA5 [19]. Therefore, we tested colocalization of pkp4 and p120 with these two proteins. Double staining clearly showed that PLEKHA5 and pkp4 were indeed colocalized in the lateral-sAJs (Fig. 2b) that was also reflected by high pkp4-PLEKHA5 pixel correlation (r ∼ 0.76, Fig. 2d). While Erbin was more evenly distributed, both visual inspection and scatterplots also showed some preference of this protein for the pkp4-enriched AJs (Fig. 2d,e). Altogether, the initial characterization of p120-CCC or pkp4-CCC suggested that they prefer different types of AJs: the pAJs (both, apical and basal location), and the lateral-sAJs, correspondingly.

### Lateral-sAJs are persistent but mobile structures

The absence of vinculin or afadin (which are important for strengthening the CCC-actin bonds) in the lateral-sAJs suggested that these junctions might be unstable. To test this idea, we performed time-lapse imaging of A431 cells in which their endogenous E-cadherin was replaced with mGFP-tagged E-cadherin (EcGFP). One-hour long movies taken at 30 sec intervals showed that the lateral-sAJs were mobile but surprisingly stable: the majority of them persisted during the entire 1h long observation (Fig. 3a,b and movie S1). Tracing their trajectories showed that the lateral-sAJs exhibited oscillatory motion that also had been reported for these junctions in other cells [14, 17]. However, we did not observe pulsatile congression of lateral-sAJs, as reported by Wu et al. [16] in Caco2 colon carcinoma cells. Instead, lateral-sAJs drifted without any specific directionality, although the neighboring junctions often showed apparently coordinated movement (see tracks shown in red in Fig. 3c). The fusion or fission of the lateral-sAJs were also often observed (Fig. 3a). The lateral-sAJs were located immediately beneath a row of relatively immobile apical-pAJs and above the area occupied by the basal-pAJs. The latter junctions, as had been reported [14, 15], were continuously generated at the basal end of the lateral plasma membranes and then moved upward (see “green” tracks in Fig 3c). The basal-pAJs were relatively short-lived since in the course of the upward movement they either merged with apical-pAJs or arrived at the zone occupied by the lateral-sAJs, where they eventually disappeared or occasionally were converted into lateral-sAJs. Thus, the lateral membranes of A431 cells comprise of three zones, each of which contain a specific type of AJs (Fig. 3c).

**Figure 3.**
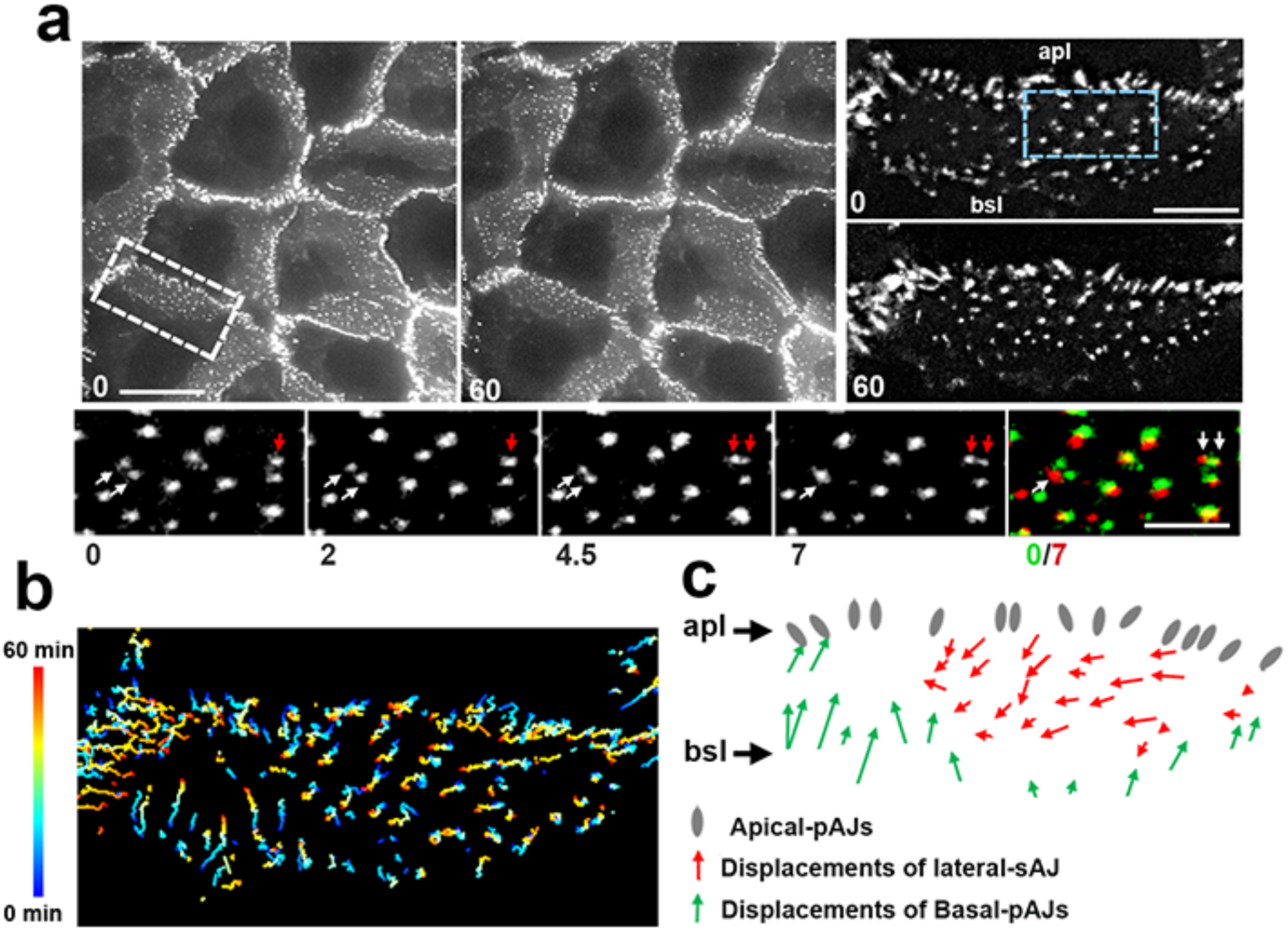
General topography and dynamics of AJs in A431 cells. (**a**) Time-lapse images of EcGFP-expressing A431 cells acquired at 30-sec intervals. Only the frames taken at the first (0) and final (60) minutes of observation are shown. Scale bar, 20 μm. The zoomed cell-cell contact (highlighted by the dashed white box) of the same frames (0 and 60 min) is shown on the right (see Movie S1). The apical (apl) and basal (bsl) edges of the contact are indicated. Scale bar, 7 μm. The time-lapse series (the bottom row) spanning 7 min of the Movie S1 demonstrates a fusion (white arrows) and a fission (red arrows) of lateral-sAJs within the area indicated by the blue dashed box (the numbers under the frames show minutes). The right image is an overlay of the initial (colorized in green) and final (colorized in red) frames of this short sequence. Scale bar, 4 μm. (**b**) The tracked trajectories of major AJs in the cell-cell contact depicted by the white dashed box in (a). The color-code of the trajectories depending on time is shown on the left. (**c**) The schematic representation of net displacements of major AJs according to tracks shown in (b). Note that the lateral membrane is clearly demarcated into three distinct zones exhibiting apical-pAJs (grey), lateral-sAJs (red), and basal-pAJs. Only approximate positioning of the apical-pAJs is shown.

### Pkp4 and p120 support distinct types of AJs

To test the role of pkp4 and p120 in AJ diversification, we knocked out pkp4 or p120 from our EcGFP-expressing cells. As expected, the p120 knockout resulted in the dramatical reduction (up to 90%) of the EcGFP expression level (Fig. 4a). No change in EcGFP expression was noted in pkp4-KO cells. However, both p120-KO and pkp4-KO cells formed AJs (Fig. S1a). Furthermore, excellent linear relationship (r ∼ 0.9) between EcGFP and pkp4 in p120-KO cells, or between EcGFP and p120 in pkp4-KO cells, strongly suggested that nearly all AJs in these cells were built from pkp4-CCC or p120-CCC respectively (Fig. S1a,b).

**Figure 4.**
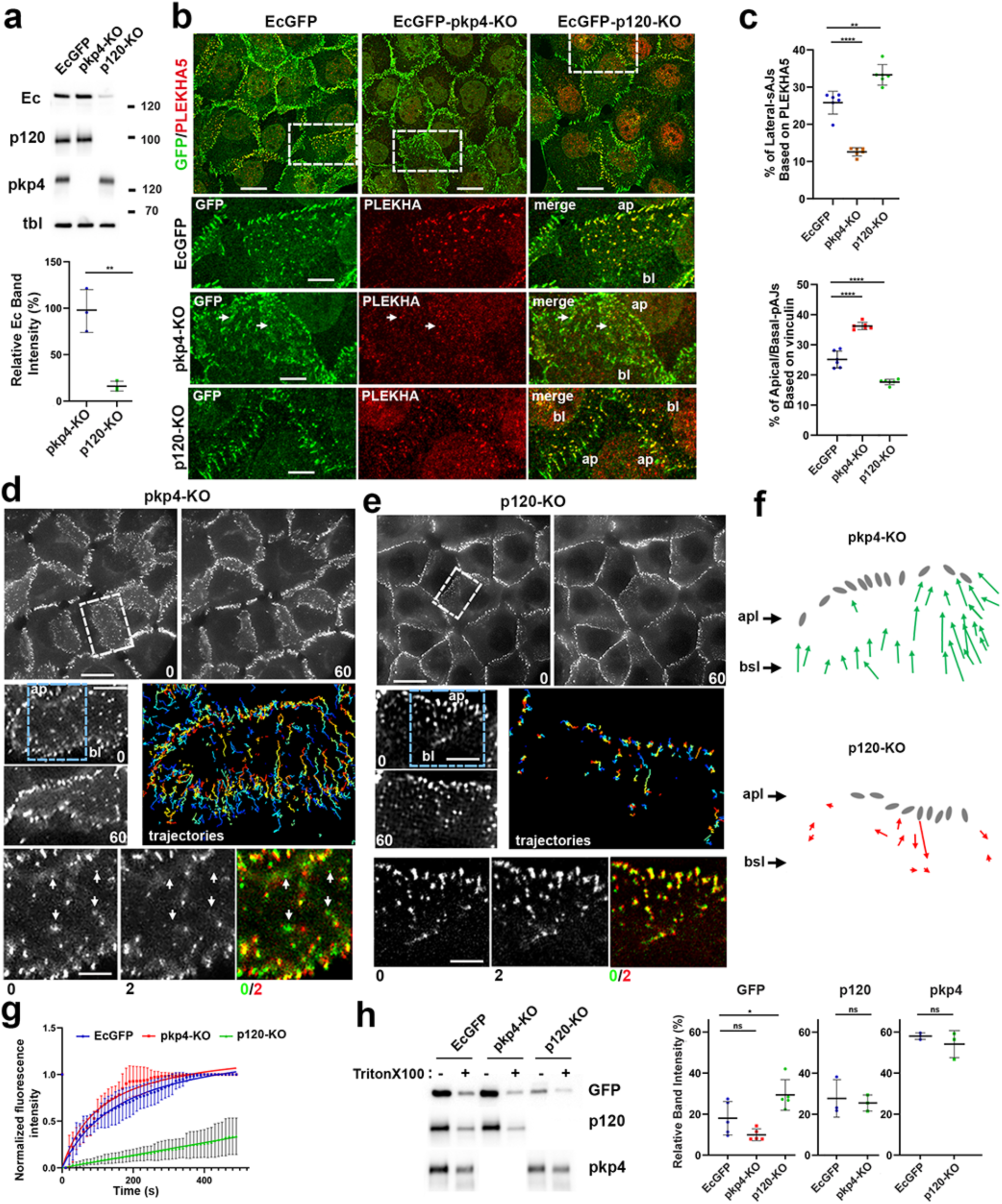
Changes in AJs upon p120 and pkp4 knockout. (**a**) Western blot of A431-EcGFP cells (EcGFP) and their pkp4-KO and p120-KO progenitors probed for EcGFP (Ec), p120, and pkp4. β-Tubulin staining (tbl) serves as a loading control. Molecular weight markers (in kDa) are shown on the right. The EcGFP band intensities in the KO cells (relatively to the parental EcGFP cells) are quantified at the bottom. (**b**) Projections of all x-y optical slices of EcGFP cells and their KO counterparts stained for EcGFP (GFP, green) and PLEKHA5 (PLEKHA, red). Only merged images are shown for low magnifications. Bar, 20 μm. The zoomed areas (marked by dashed boxes) are shown in both colors at the bottom (ap and bt indicate apex and bottom of the contacts). Bar, 12 μm. The arrows point at few remaining PLEKHA5-positive AJs in p120-KO cells. (**c**) Amounts of heterochromatic EcGFP/PLEKHA5-positive (derived from lateral-sAJs) or EcGFP/vinculin-positive (derived from apical/basal-pAJs) pixels relative to the total number of EcGFP-positive pixels (n = 6 taken from the three independent images). The means +/-SD are indicated by bars. (**d, e**) Time-lapse images of EcGFP-pkp4-KO (**d**) and EcGFP-p120-KO (**e**) cells acquired at 30-sec intervals. Only the first (0) and final (60) frames are shown (see Fig. 3 for details). Scale bar, 20 μm. The zoomed cell-cell contacts (indicated by dashed white boxes) from each of the sequences are shown at the bottom (see also Movie S2 and S3). The apical (ap) and basal (bl) edges of the contacts are indicated. Scale bar, 7 μm. The zoomed areas (indicated by blue dashed boxes) of these contacts taken 2 minutes apart presented at the bottom row demonstrate instabilities of the lateral AJs in pkp4-KO cells and their stability in p120-KO cells (the numbers under the frames show minutes). The right image is an overlay of the first (colorized in green) and the second (colorized in red) frames. The arrows point to the AJs that are disassembled over this time. Scale bar, 4 μm. The tracked trajectories of major AJs (trajectories) are shown as in Fig. 3b. (**f**) The schematic representation of net displacements of major AJs according to the trajectories shown in **d** and **e**. The AJs are depicted as in Fig. 3c. (**g**) FRAP assay of lateral-sAJs in cells indicated as in a (n = 15). (mean +/− SEM). (**h**) Left: Western blot probed for EcGFP, p120, and pkp4 of total cell lysates (cells are indicated as in a) from control culture (-) and from parallel cultures after 5 min-long extraction with 1% Triton X100 (+). Right: Quantification (based on five independent experiments) of the intensities of EcGFP relative to that in nonextracted cultures. Statistical significance for all graphs was calculated using a two tailed Student’s t test: ns, non-significant; *, P < 0.05; **, P<0.01; ***, P< 0.001; ****, P < 0.0001. The means +/-SD are indicated by bars.

To elucidate the types of AJs in cells lacking p120 or pkp4, the cells were stained for vinculin and PLEKHA5, best available markers for apical/basal-pAJs and lateral-sAJ, respectively (Fig. 4b and S1c). It showed that both types of knockout cells still produced both types of AJs. However, their abundance was knockout-specific. Many apical AJs and nearly all basally located AJs in the p120-KO cells exhibited almost no vinculin staining (Fig. S1c) but were positive for PLEKHA5 (Fig. 4b), suggesting a predominance of the lateral-sAJs in these cells. By contrast, PLEKHA5 in AJs of pkp4-KO cells was detected only occasionally (Fig. 4b). Vinculin in these cells was detected even in some AJs located at the middle portion of the lateral membrane (Fig. S1c). Such a disbalance between the lateral-sAJs and apical/basal-pAJs in the KO cells was validated by quantification of the relative abundance of these junctions. To estimate this parameter, we determined the ratio of the heterochromatic pixels, positive for E-cadherin/PLEKHA5 (derived from lateral-sAJs) or E-cadherin/vinculin (derived from pAJs) versus the total number of the E-cadherin-positive pixels (derived from all AJs). It confirmed that the abundance of the lateral-sAJs dropped by ∼ 70% in the pkp4-KO cells but increased by ∼ 50% in the p120-KO cells, as compared with the parental EcGFP-expressing cells. In contrast, the fraction of apical/basal-pAJs was increased upon pkp4 depletion and decreased in absence of p120 (Fig. 4c).

The time-lapse imaging revealed even more dramatic differences between p120-KO and pkp4-KO cells. Majority of the AJs in pkp4-KO cells showed the upward movement typical for basal-pAJs (Fig. 4d,f and Movie S2). In addition, these cells showed high instability of AJs located at the lateral surface: each lateral membrane continuously displayed their rapid disappearance. This process usually started with conversion of the lateral AJs into a small fluorescent “cloud”. Figure 4d (the bottom row) highlights four such events occurring in just two initial minutes of the 60-min long movie (between frames 0 and 4 of the Movie S2) within a small area of the lateral membrane. In contrast to pkp4-KO cells, no steady upward movement of AJs was detectable in the p120-KO cells (Fig. 4e,f and Movie S3). Also, these cells, like the parental EcGFP cells, showed long-term persistence of their lateral AJs lasting during the entire observation (see the bottom row in Fig. 4e). Taken together, these data suggested that pkp4-CCC and p120-CCC, while able to form all types of AJs, are better designed for lateral-sAJs and apical/basal-pAJs, correspondingly.

### Pkp4 and p120 maintain the opposite AJ dynamics

One explanation for these results is that p120, but not pkp4, maintains fast CCC turnover that destabilizes the lateral-sAJs concurrently promoting directional movement of the basal-pAJs. To test this idea, we performed Fluorescence Recovery after Photobleaching assay (FRAPA). The individual AJs located at the lateral membrane were photobleached and their fluorescence recovery was traced over about 6 min (Fig. 4g). Indeed, the p120-deficient AJs were exceptionally stable: they recovered only about 20% of their fluorescence during the observation period. The pkp4-deficient junctions were, by contrast, the most dynamic. They reached nearly 100% recovery in about 5 min showing the halftime of recovery (t1/2) of about 1 min. The wt lateral-sAJs showed the intermediate phenotype (t1/2∼2 min).

In parallel experiments we elucidated the anchorage of p120-CCC and pkp4-CCC to the cytoskeleton by testing their resistance to the Triton-X100 treatment. To this end, the levels of E-cadherin, p120, and pkp4 remaining in cells after Triton X100 extraction were compared with those in the untreated cells. The results showed that ∼ 20% of EcGFP and p120 remained bound to the control and pkp4-KO cells after Triton extraction (Fig. 4h). This EcGFP fraction was slightly higher (up to 30%) in the p120-KO cells. Importantly, the Triton X100-resistant fraction of pkp4 was significantly higher (∼ 60%) in both control and p120-KO cells (Fig. 4h). Assuming that the level of pkp4 reflects the level of pkp4-CCC, these results indicate that this complex is more tightly bound to the cytoskeleton than p120-CCC.

### Recombinant pkp4 and p120 facilitate formation of lateral-sAJs and pAJs, correspondingly

The differences in AJ phenotypes between control, pkp4-KO and p120-KO cells suggested a role of pkp4 in reinforcement of the lateral-sAJs. However, the abundance of highly stable lateral-sAJ in p120-KO cells could be also explained by low EcGFP level in these cells (see Fig. 4a) that could indirectly perturb the cadherin turnover in AJs. ARVCF, the third δ-catenin of A431 cells, also complicates the interpretation of our results. To convincingly show that pkp4 and p120 play specific roles in formation and/or maintenance of distinct types of AJs, we generated A431 cells lacking all three endogenous δ-catenins (p120, pkp4, ARVCF) and then reexpressed in the obtained δ-catenin null cells (A431-δCat-KO cells) the GFP-tagged versions of pkp4 (GFPpkp4) or p120 (GFPp120). The Western blot characterization of the obtained cells (Fig. 5a) showed that the A431-δCat-KO cells exhibited only traces of E-cadherin, but as expected, its expression was increased (while not reaching the wt level) upon GFPpkp4 transfection. Noteworthy, staining of the blots for pkp4 showed that GFPpkp4 was dramatically overexpressed in the GFPpkp4-δCat-KO cells compared with the endogenous pkp4 in the wt A431 cells. However, based on GFP staining, this GFPpkp4 level was still significantly lower than the GFPp120 level in GFPp120-δCat-KO cells (Fig. 5a).

**Figure 5.**
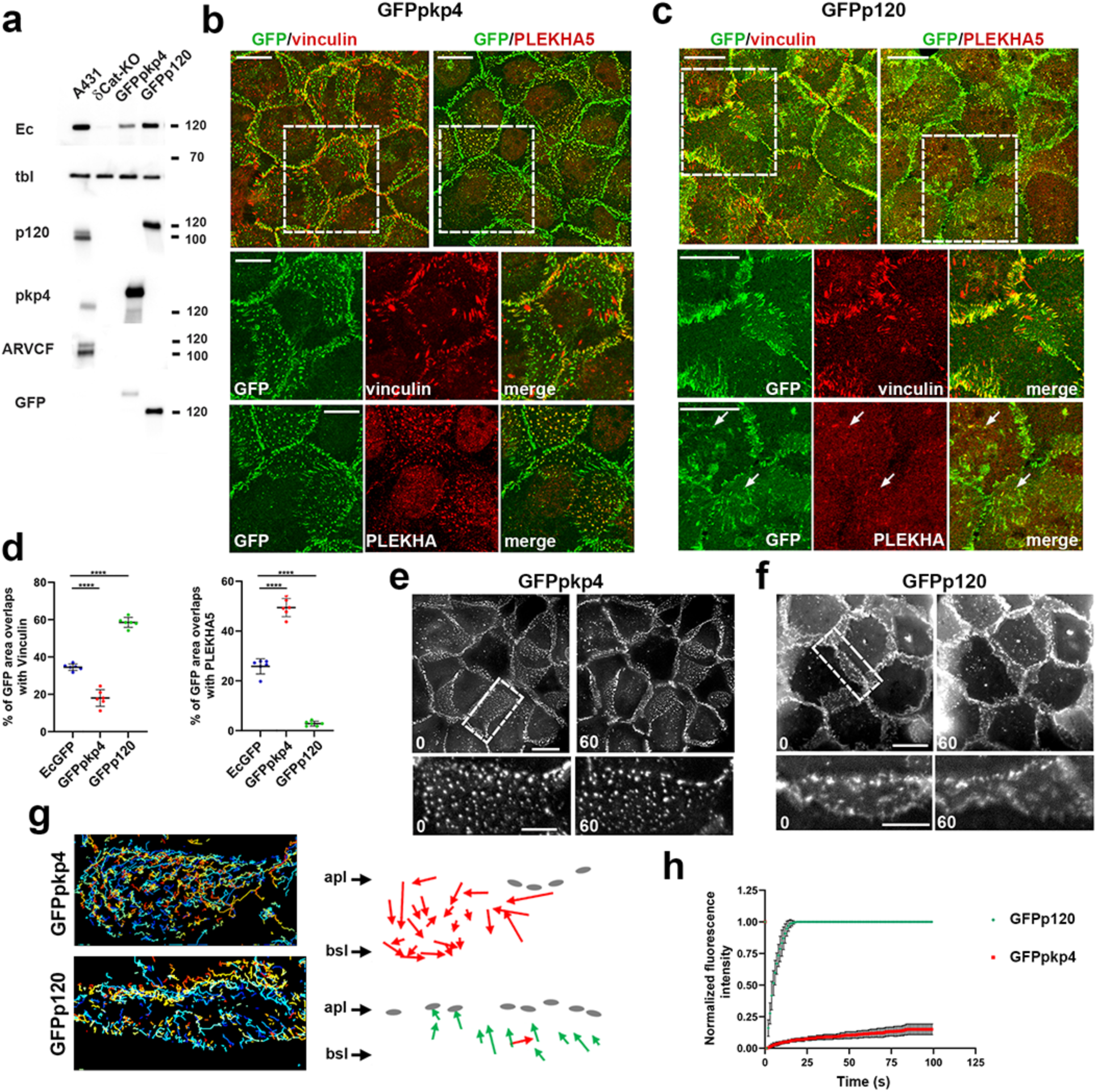
GFPpkp4 and GFPp120 promote formation of distinct types of AJs. (**a**) Western blot of wt A431 cells (A431), δCat-KO counterparts, and their GFPpkp4 and GFPp120 progenitors probed for E-cadherin (Ec), p120, pkp4, ARVCF, and β-tubulin (tbl) as a loading control. Molecular weight markers are shown as in Fig. 4a. Note that both GFPp120 and GFPpkp4 expression increase the level of E-cadherin. Note also that GFPpkp4 is significantly overexpressed while still remains below the endogenous level of p120 (compared endogenous and recombinant proteins in pkp4 and p120 staining). (**b, c**) Projections of all x-y optical slices of GFPpkp4 (**b**) and GFPp120 (**c**) cells stained for GFP (green) and AJ type markers, vinculin and PLEKHA5 (red). Only merged images are shown for low magnifications. Bar, 20 μm. The zoomed areas (marked by dashed boxes) are shown in both colors at the bottom. Bar, 15 μm. The arrows point at some of the remaining PLEKHA5-positive AJs in p120-KO cells. (**d**) Amounts of heterochromatic GFP/PLEKHA5-positive (derived from lateral-sAJs) or GFP/vinculin-positive (derived from apical/basal-pAJs) pixels relative to the total number of GFP-positive pixels (n = 6 taken from the three independent images). Statistical significance was calculated using a two tailed Student’s t test: ****, P < 0.0001. The means +/-SD are indicated by bars. Note that GFPp120 and GFPpkp4 promote opposite effects on amounts of the sAJ and pAJs. (**e, f**) Time-lapse images of cells expressing GFPpkp4 (**e**) and GFPp120 (**f**) acquired at 30-sec intervals. Only the first (0) and final (60) frames are shown. Scale bar, 20 μm. The zoomed cell-cell contacts (indicated by dashed white boxes) from each of the sequences are shown at the bottom (see also Movie S4 and S5). The apical (ap) and basal (bl) edges of the contacts are indicated. Scale bar, 10 μm. Note a big difference in overall organization of AJs between the cell lines. (**g**) The trajectories of major AJs are shown as in Fig. 3b at the left. At the right side, the schematic representation of net displacements of major AJs according to these trajectories. The AJs are depicted as in Fig. 3c. (h) FRAP assay of lateral-sAJs in cells indicated as in a (n = 15). (mean +/− SEM).

In line with the data reported above for the EcGFP-expressing p120-KO cells, the GFPpkp4-expressing δCat-KO cells produced numerous prominent lateral-sAJs (Fig. 5b and S2). Nearly perfect co-localization of GFPpkp4 with E-cadherin (r ∼ 0.9) also confirmed that all AJs in these cells were made of GFPpkp4-CCC (Fig. S2). Also, in agreement with the results obtained for p120-KO cells, all lateral-sAJs and even some of AJs formed at the basal or apical edges of the lateral membrane (e.g., at sites of basal-or apical-pAJs) showed no vinculin staining (Fig. 5b). Accordingly, quantification of the heterochromatic vinculin/GFP and PLEKHA5/GFP pixels as readouts for two types of AJs revealed that GFPpkp4-expressing δCat-KO cells, similar to the p120-KO cells and in contrast to the control EcGFP-expressing cells, produced an increased pool of lateral-sAJs at the expense of the apical/basal-pAJs (Fig. 5b,d). The prevalence of the lateral-sAJs in these cells was further confirmed by time-lapse imaging (Movie S4) and FRAP experiments: AJs in GFPpkp4-expressing cells showed no directional upward movement (Fig. 5e,g) coupled with extremely low rate of fluorescence recovery (Fig. 5h). Thus, the cells expressing GFPpkp4 and deficient for other δ-catenins overproduced lateral-sAJs.

The analogous experiments with GFPp120-expressing cells showed that they formed AJs which phenotype matched that of AJs in pkp4-KO cells. Specifically, these cells showed a dramatic drop in the PLEKHA5/GFP colocalization concurrently with an increase in vinculin/GFP heterochromatic pixels (Fig. 5c,d). The nearly complete absence of lateral-sAJs was further corroborated by time-lapse microscopy. It showed that GFPp120-expressing cells predominantly produced upward moving short-lived AJs, which often disintegrated through an intermediate “cloud-like” transformation (Movie S5, Fig. 5f,g,h). Accordingly, FRAP assay showed dramatic difference between the rates of GFPp120 and GFPpkp4 turnover (Fig. 5h). Collectively, these experiments showed that p120 and pkp4 have a great impact on global organization of the cell-cell adhesion system and the types of AJs.

### Pkp4-CCC clusters are formed along actin filaments in α-catenin independent manner

One distinguishing feature of the lateral-sAJs is the lack of vinculin and afadin. These proteins associate with the CCC through tension-dependent changes in the α-catenin conformation [32–35]. Why then, do lateral-sAJs, despite their exceptional strength coupled with apparently actomyosin-dependent oscillatory movements [16, 17], lack these two tension-dependent proteins? To explain this paradox, we proposed that pkp4-CCC might interact with actin filaments using an α-catenin-independent mechanism. To test this possibility, we used the α-catenin-deficient αCat-KO-A431 cells expressing an mCherry-tagged α-catenin mutant, αCatCH-Δ259. Its extensive deletion (aa 260-906) removed all known α-catenin determinants directly or indirectly connecting α-catenin to F-actin while preserving its N-terminal β-catenin-binding domain (Fig. 6a).

**Figure 6.**
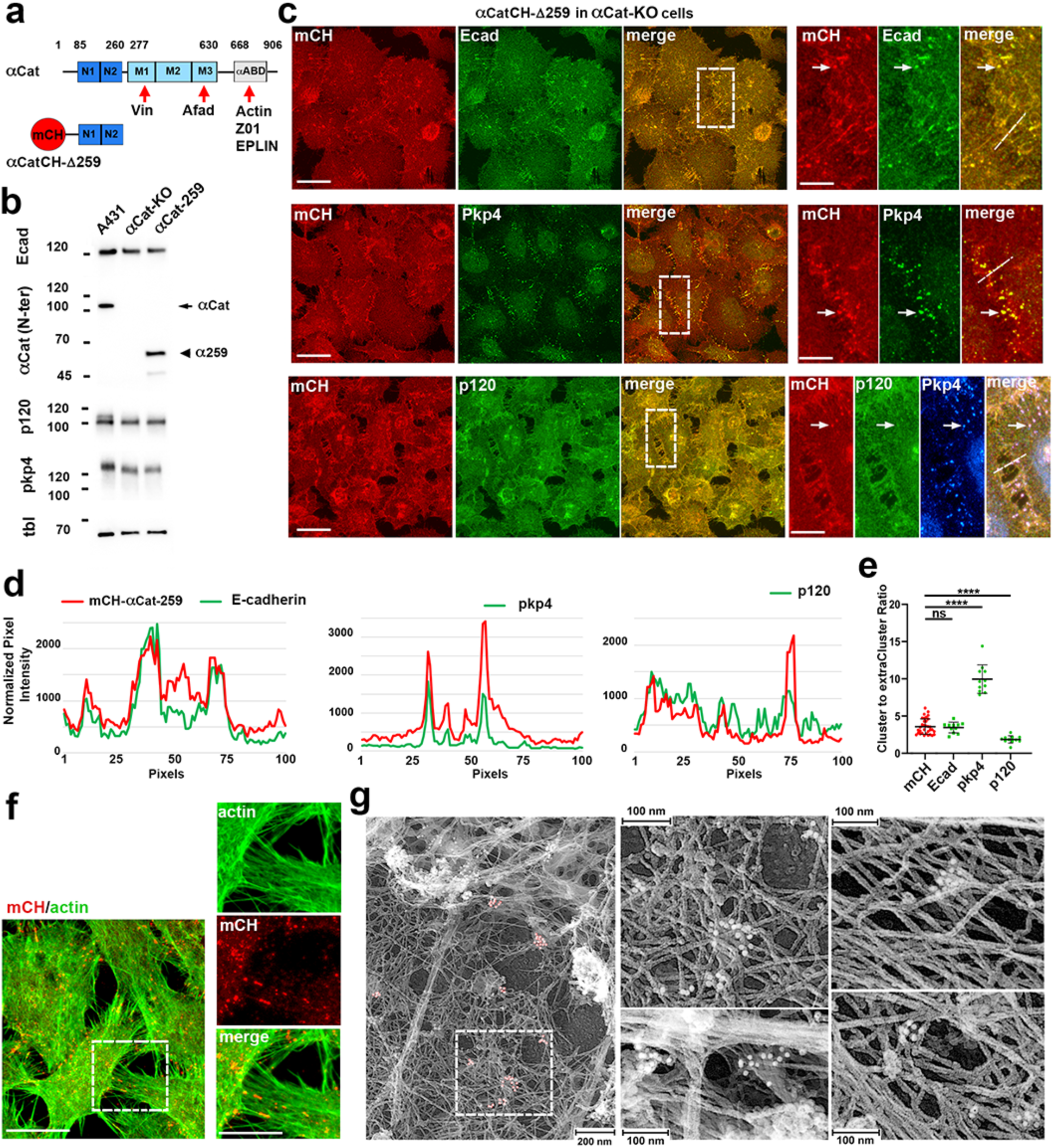
Pkp4 facilitates formation of the α-catenin-independent clusters associated with actin filaments. (**a**) Schematic representation of the intact α-catenin (αCat) and the α-catenin mutant, αCatCH-Δ259 consisting of only the β-catenin-binding N-terminal α-catenin region (residues 1-259). All known sites that couple α-catenin to F-actin or to the actin-associated proteins, such as vinculin (Vin), afadin (Afad), ZO1, EPLIN, which all are in the M subdomains (M1, M2, M3) and in the actin-binding domain (αABD), are deleted. Red circle (mCH) depicts mCherry, the unstructured regions are shown as solid lines. The borders between domains are indicated by the numbers of corresponding residues. (**b**) Western blot of wt A431 cells (A431), αCatKO-A431 cells (αCat-KO) and the αCat-KO cells expressing αCatCH-Δ259 (αCat-259) probed for E-cadherin (Ecad), N-domain of α-catenin (αCat, N-term), p120, and pkp4 and β-tubulin (tbl) as a loading control (tbl). Molecular weight markers (in kDa) are on the left. The intact α-catenin and the mutant are indicated on the left. (**c**) Projections of all x-y optical slices of αCatCH-Δ259 cells double stained for mCH (red) and E-cadherin (Ecad) or pkp4 (green), or triple stained for mCH, p120, and pkp4 (blue, shown only for the zoomed area). Bar, 20 μm. The zoomed areas (marked by dashed boxes) are shown at the right side. Bar, 6 μm. The arrow points at one of the clusters in each zoomed image. (**d**) The line scan performed along the dashed line shown in (c). (**e**) The ratio between cluster and inter-clusters mCH, E-cadherin (Ecad), pkp4, and p120fluorescence. (n = 11 taken from two independent images). The means +/-SD are indicated by bars. (**f**) Projections of all x-y optical slices of αCatCH-Δ259 cells extracted with 1% Triton-X100 and stained for mCH (red) and F-actin (green). Bar, 20 μm. The zoomed areas (marked by dashed boxes) are shown at the right side. Bar, 10 μm. (**g**) PREM of αCatCH-Δ259 clusters. The left image shows an overview of two overlapped cells. The boxed regions are enlarged in the middle (top). Other images show other examples of the clusters taken from other cells in the same culture. mCH immunogold particles (10 nm, red) are pseudocolored.

Western blotting showed that the cells transfected with αCatCH-Δ259 expressed the same levels of E-cadherin, p120, and pkp4 as wt A431 cells and an expression level of the mutant matched that of the endogenous α-catenin (Fig. 6b). As expected, the majority of the mutant, as well as E-cadherin, were diffusely distributed along the entire plasma membrane of these cells (Fig. 6c). However, in addition to this broad distribution, the arrays of small clusters incorporating both proteins could be noticed along the filopodia-like protrusions connecting the cells. Remarkably, pkp4 specifically resided in these clusters (Fig. 6c,d). The recruitment of pkp4 into these clusters appeared to be very efficient since, by contrast to the αCatCH-Δ259 mutant or E-cadherin, little if any pkp4 was detected outside the clusters (see line scans in Fig. 6d). This observation was further confirmed by evaluating the ratio between fluorescence within versus outside of the clusters: while the fluorescence intensity of E-cadherin or αCatCH-Δ259 in the clusters exceeded the surrounding areas just about four times, this parameter was ∼10 for pkp4 (Fig. 6e). The cluster/extra-cluster ratio was even less for p120 (∼ 3), and this protein was hardly detectable in some of the clusters (Fig. 6c-e).

We then examined a possibility that the αCatCH-Δ259 clusters were caused by vinculin, which in some cells can directly interact with β-catenin [36–38]. Staining the cells for vinculin, however, showed no association of this protein with the αCatCH-Δ259/pkp4 clusters (Fig. S3a).

Finally, stained for erbin and PLEKHA5 (Fig. S3b) verified that by protein composition the αCatCH-Δ259/pkp4 clusters did correspond to the lateral-sAJs. Altogether, these data provide strong evidence that the pkp4-CCC generates clusters through a specific mechanism that bypasses α-catenin-F-actin interactions.

Formation of the αCatCH-Δ259/pkp4 clusters along the actin-rich cell-cell protrusions suggested that despite of α-catenin independence, the pkp4-CCC clusters still use actin filaments for their formation. To validate this point, we removed most of the free cytosolic proteins by gentle extraction of the cells with 1% Triton X100 in cytoskeleton preservation buffer, and then the Triton-resistant cytoskeleton was stained for mCherry and F-actin. This staining showed that the αCatCH-Δ259 clusters were Triton X100-resistant. The brightest clusters were mostly observed along the actin bundles associated with the cell protrusions. In addition, numerous smaller clusters were dispersed along the actin cortex of the overlapping lamellipodia of the adjacent cells (Fig. 6f). To provide better understanding of the relationship between the clusters and F-actin, we then performed platinum replica electron microscopy (PREM) in combination with αCatCH-Δ259 immunogold labeling of the Triton-X100 extracted cells. In these experiments, we concentrated on the anti-mCherry immunogold within well spread areas of overlapped actin networks of the neighboring cells, where both the gold particles and individual actin filaments could be easily identified. Strikingly, the immunogold particles were organized in numerous clusters of different sizes, all of which were clearly associated with actin filaments especially often in the sites where several filaments merged into multifilament sheets (Fig. 6g).

Altogether, these experiments showed that pkp4-CCC form actin-associated clusters bypassing the α-catenin-dependent mechanism participating in formation of the tension-dependent AJs.

### Pkp4-faclitated CCC clustering has no role in cell-cell adhesion

We next sought to understand whether the α-catenin-independent pkp4-CCC clusters contribute to cell-cell adhesion. To study this question, we used the dispase adhesion assay to compare control EcGFP cells with their pkp4-KO and p120-KO counterparts, in which the pkp4-or p120-facilitated clustering were abolished. As expected, after dispase-mediated detachment of cells from the substrate, the confluent monolayers of the control EcGFP-expressing cells contracted due to the contractile forces within the epithelial sheet. This contraction, quantified based on the sheet areas, was not significantly perturbed in pkp4-depleted cells (Fig. 7a). The cell-cell adhesion strength in these cells was only slightly weakened, as was evident from the small but statistically significant increase in the number of fragments generated by mechanical stress applied to the lifted sheets. Thus, the shortage of lateral-sAJs does not significantly interfere with general cell-cell adhesion. As expected, both cell sheet contraction and adhesion strength were severely affected in p120-KO cells (Fig. 7a). Most likely, these dramatic effects observed in p120-KO cells were caused by the reduced level of E-cadherin but not by general inability of pkp4-CCC to maintain cell-cell adhesion.

**Figure 7.**
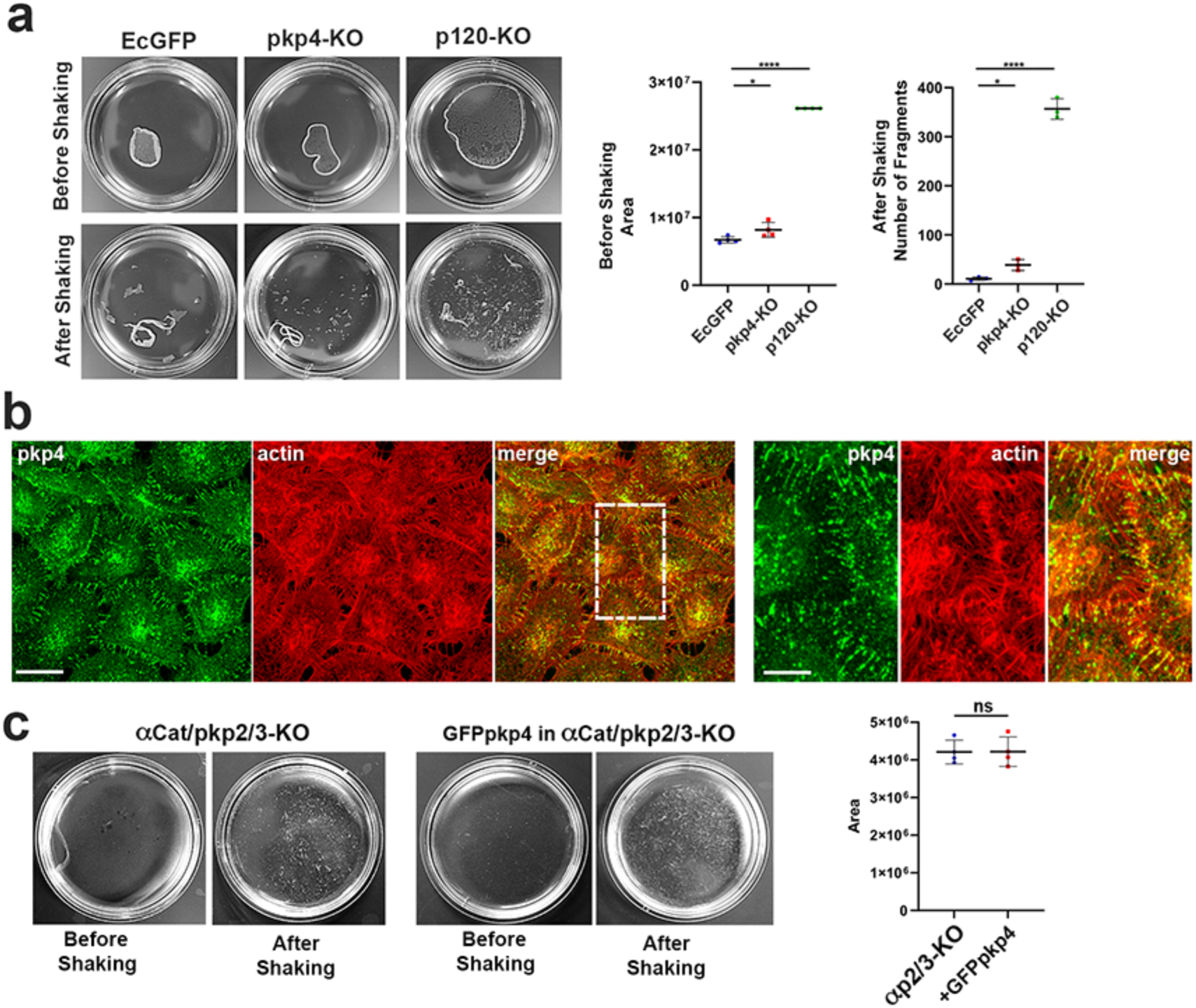
Pkp4-facilitated E-cadherin clusters have no adhesion function. (**a**) Dispase-based assay of A431 cells expressing EcGFP (EcGFP) and their progenitors deficient for pkp4 (pkp4-KO) or for p120 (p120-KO). Left: Representative images showing the cell sheets detached from the dishes before and after mechanical stress (shaking). Right: Quantification (n = 4) of the areas (in pixels) of the sheets before shaking (left) and the number of fragments obtained after shaking (right). Data are presented as mean values +/-SD. (**b**) Projections of all x-y optical slices of αCat/pkp2/3-KO cells expressing recombinant GFPpkp4 stained for pkp4 (green) and F-actin (red). Bar, 20 μm. The zoomed area (marked by white dashed boxes) is shown at the right panel. Bar, 8 μm. Note a perfect alignment of pkp4 clusters along the actin-rich cell-cell protrusions. (**c**) Dispase-based assay of the parental αCat/pkp2/3-KO cells (αCat/pkp2/3-KO) and their counterparts expressing GFPpkp4 (GFPpkp4-αCat/pkp2/3-KO). Left: Representative images showing the cell sheets detached from the dishes before and after mechanical stress (shaking). Note that the cell sheets of both cell lines are completely unable to contract. Right: Quantification (n = 4) of the areas (in pixels) of the sheets before shaking (left). The number of fragments obtained after shaking was not quantified. Data are presented as mean values +/-SD. Statistical significance was calculated using a two tailed Student’s t-test: ns, non-significant; *P < 0.05; **P < 0.01; ***P < 0.001; ****P < 0.0001.

To directly test whether α-catenin-independent pkp4-CCC clusters can maintain cell-cell adhesion, we generated αCat/pkp2/3-KO A431 cells and their counterparts overexpressing GFPpkp4. In addition to the lack of α-catenin, the αCat/pkp2/3-KO A431 cells were also deficient for two desmosomal plakophilins, pkp2 and pkp3. These two desmosomal pkp4 relatives were shown to be essential for desmosome (DSM) assembly [39–41]. Indeed, staining for Dsg2 showed that αCat/pkp2/3-KO A431 cells were unable to form DSMs (Fig. S4). The absence of DSMs excluded the contribution of DSMs to cell-cell adhesion thus allowing to detect even weak increase in adhesion upon GFPpkp4 expression. Despite a complete absence of α-catenin and DSMs, the endogenous pkp4 in the αCat/pkp2/3-KO cells as well as recombinant GFPpkp4 in their GFPpkp4-expressing variant still efficiently produced clusters, which were predominantly located along the actin-rich intercellular protrusions (Fig. S4 and 7b). However, the pkp4-CCC clusters even in GFPpkp4-expressing cells were completely unable to mediate any contraction forces as well as to maintain measurable intercellular adhesion (Fig. 7c).

## Discussion

Cadherin-mediated adhesions are operated by two CCC oligomerization processes taking place outside and inside the cells. The “outside” cadherin oligomerization forms an adhesive paracrystalline lattice where extracellular cadherin regions are coordinated by a combination of *trans* and *cis* bonds [1, 42–44]. The “inside” CCC oligomerization is mediated by α-catenin-actin interactions and generates linear actin-bound CCC strands [45–47]. Working synergistically, these two relatively independent “outside” and “inside” oligomerization processes produce CCC clusters, which adhesion strength could be tuned by cell signaling mechanisms targeting the α-catenin-F-actin association. Understanding this regulation is certainly a major direction for future research. However, our current knowledge of CCC clustering in AJs is apparently incomplete.

Available data suggest that the two oligomerization processes described above could be only a fraction of the versatile binding cascades that eventually produce amazing diversity of AJs that have been demonstrated in cells [9, 10]. For example, the general structure of invertebrate and vertebrate cadherins shows that they form different “outside” oligomers [48]. A novel type of cadherin ectodomain dimerization has been revealed in a recent Cryo-EM study [49]. It is also clear that α-catenin interactions with F-actin, depending on the involvement of additional adaptor proteins, generate supramolecular structures of different strength and architecture [50, 51]. Finally, junction-associated cadherin clusters were detected in α-catenin-deficient cells [52–54]. Our study reported here provides strong evidence for the alternative pkp4-dependent “inside” CCC oligomerization, which plays a key role in AJ diversification.

Using two epithelial cell models, A431epidermal carcinoma and DLD1 colon carcinoma, we show that pkp4, contrary to more widespread δ-catenin, p120, preferably incorporates into the lateral-sAJs. These AJs are known to reside in the middle domain of the lateral membrane in nearly all epithelial cells. They can be distinguished from more apically and more basally located AJs by three features: (i) deficiency for vinculin and afadin [12]; (ii) abundance of PLEKHA5 and erbin, though they both, but especially the latter, could be found in isolated clusters within other types of AJs [18, 19]; and (iii) their nondirectional oscillatory motion [16, 17]. By tracking these junctions in A431 cells we also found that despite being constantly in motion, they are incredibly durable with respect of their lifetime.

Our examination of pkp4-KO cells shows that despite a preferential recruitment of pkp4 into lateral-sAJs, pkp4 is not essential for this type of junctions. Nevertheless, the pkp4 depletion greatly reduces the number of lateral-sAJs. Furthermore, most of the AJs retained at the lateral membranes in pkp4-KO cells appear to belong to the basal-pAJ category. The phenotype of pkp4-KO cells is limited to AJs located at the lateral membrane, since these cells do not demonstrate obvious defects in apical/basal-pAJs, nor they show significant abnormalities in cell-cell adhesion evaluated by the dispase assay. A role of pkp4 in the maintenance of the lateral-sAJs is also highlighted by the fact that p120 depletion that elevates the relative level of pkp4-CCC results into the exactly opposite outcome: majority of AJs in p120-KO cells bear the traits of the lateral-sAJ. Such instructive role of pkp4 in AJ specialization is further confirmed by overexpression of the recombinant form of pkp4 in δ-catenin-null cells. These GFPpkp4-expressing cells, while still able to form pAJs marked by vinculin, show significant and specific overproduction of the lateral-sAJs. Remarkably, the same δ-catenin-null cells expressing GFPp120, by contrast, overproduce apical-and basal-pAJs but show a clear shortage of lateral-sAJs.

Altogether, our comparative study of two δ-catenins, p120 and pkp4, using both loss and gain of function experiments, clearly shows that p120 is designed for tension-dependent AJs, such as apical or basal-pAJs, where “intracellular” clustering is based on the canonical α-catenin-actin interactions. By contrast, pkp4 supports the strength of the lateral-sAJs. The absence of vinculin and afadin in the latter junctions suggests that the remarkable strength of the lateral-sAJs is not based on the vinculin/afadin-mediated reinforcement of α-catenin-actin bonds. Our interrogation of the canonical α-catenin-actin interactions indeed shows that the canonical α-catenin-actin interactions are suppressed in the lateral-sAJs. Instead, to reinforce the “outside” cadherin adhesive lattice these junctions use an alternative “inside” clustering mechanism.

According to previous data [47], A431 cells lacking functional α-catenin produce no measurable cell-cell adhesion in the adhesion assay. Nor they form AJs anchored to the radial actin bundles. However, the confocal microscopy in conjunction with immunogold PREM shows that these cells assemble pkp4-CCC clusters, which are associated with the cortical actin cytoskeleton. At the light microscopy level, these clusters recruit PLEKHA5 and erbin that underscores their lateral-sAJ identity. Another feature of these α-catenin-independent clusters is that they exhibit only negligible incorporation of p120. A stark difference in subcellular localization of p120-CCC and pkp4-CCC is a clear indication of differences in the clustering mechanisms of these two complexes and in their binding to actin filaments, in particular. The specific mode of pkp4-CCC interactions with actin is independently revealed by the different solubility of p120 and pkp4 upon Triton-X100 extraction in the control, α-catenin expressing, cells. Altogether, our data show that the lateral-sAJs are specialized structures that use a unique CCC clustering mechanism that includes a specific, α-catenin-independent mode of interactions with F-actin. More experiments are needed to determine whether pkp4 is directly involved in formation of α-catenin-independent CCC clusters or just “instruct” other CCC-associated proteins to produce the clusters. The first possibility is supported by our recent observation that the ARM domain of pkp3, that is the most conserved portion of the 9 ARM repeat protein family, directly interacts with actin filaments [55]. Therefore, our current working hypothesis is that pkp4-CCC, once it incorporates into the lateral AJs, activates a specific mode of interaction with actin filaments that results in pkp4-dependent CCC oligomerization.

The relationship between lateral-sAJs and DSMs is another interesting aspect for discussion. Indeed, these two evolutionary related types of junctions resemble each other by their spot-like morphology, subcellular localization, and compositions. The N-terminal portion of pkp4 was shown to interact with DSP proteins [56, 57]. Like lateral-sAJs, DSMs show exceptional stability and oscillatory motion in living cells [58, 59]. Our finding that pkp4 regulates formation of lateral-sAJs adds another similarity between these structures and DSMs.

Indeed, like pkp4 in lateral-sAJs, DSM-specific plakophilins, pkp2 and pkp3, were shown to play a key role in desmosomal cadherin clustering [39–41]. Therefore, a more complete understanding of how pkp4 facilitates production of lateral-sAJs may shed light on a long-standing question about mechanisms of DSM assembly.

In conclusion, we show here that the two distinct members of the δ-catenin protein family promote formation of two different types of AJs. The more abundant δ-catenin in cells, p120, directs formation of cadherin clusters through a clustering mechanism that includes canonical α-catenin-dependent interactions with actin filaments. This relatively well studied mode of interactions results in formation of punctate or linear AJs that are known to play a major role in cell-cell adhesion and interconnect the AJs to the actomyosin tensile forces shaping individual cells in tissues. A minor member of the δ-catenin family, pkp4, facilitates formation of the lateral-sAJs, and the underlying mechanism of their formation is an α-catenin independent. This type of junctions appears to play no role in cell-cell adhesion processes but may be important for cell-cell signal transductions since they keep lateral membranes of adjacent cells in proximity.

Collectively, out study shows a previously unrecognized function of δ-catenins -their role in regulation of the AJ assembly pathways that controls a balance between different types of AJs in cells and eventually a global organization of cell-cell adhesion system.

## Materials and Methods

### Plasmids

The plasmids (all in pRcCMV) encoding GFP-tagged pkp4 was constructed using a cDNA kindly provided by Drs Mechthild Hatzfeld (Martin Luther Univerity Halle-Wittenberg, Germany) and Kathleen J Green (Northwestern University, Chicago, USA). The plasmid encoding GFPp120 was obtained using previously published plasmid encoding p120-3A isoform of human p120 [60]. The mCherry-tagged mutant of human αE-catenin (αCatCH-Δ259) was constructed using PCR-based mutagenesis of the plasmid encoding the intact protein described previously [47]. The general map of the mutant is presented in Fig. 6a. All plasmid inserts were verified by sequencing.

### Cell culture and transfection

The original DLD1, A431 cells, EcGFP-expressing E-cadherin-deficient A431 cells (EcGFP-EcKO-A431 cells), their p120-KO progenitors, and α-catenin deficient αCatKO-A431 cells have been previously described [20, 47, 61]. The pkp4-KO version of the EcGFP-EcKO-A431 cells were obtained using the Alt-R CRISPR-Cas9 System (IDT), which had been used in our laboratory to obtain Ec-KO or p120-KO cells [20, 61]. The same strategy was used to obtain δCat-KO-A431 cells. In brief, the cells were transfected with an RNA complex consisting of a gene-specific CRISPR RNA (crRNA; designed by software of the Broad Institute of Harvard and the Massachusetts Institute of Technology) and transactivating RNA. The following crRNAs were used: pkp4-5’-AGGATCAACTAACAACCATG and ARVCF-5’-CATCCGAAGATGGCACAACC. The GFPpkp4-, GFPp120-, and αCatCH-Δ259-expressing cells were obtained using stable transfection of the corresponding cells with the corresponding plasmids. The cells were grown in DMEM supplemented with 10% FBS and were transfected using Lipofectamine 2000 (Invitrogen) according to the company protocol. After selection of the Geneticin-resistant cells (0.5 mg/ml), the cells were sorted for transgene expression by FACS, and only moderate-expressing cells were used. At least three clones were selected for each construct, and all were tested in most of the assays. The expression levels and sizes of the recombinant proteins in the obtained clones were analyzed by Western blotting as previously described [47]. All clones of cells expressing a particular transgene exhibited the same phenotype. Representative data for one of three clones is presented.

The Dispase assay was performed as described in [62]. In brief, confluent cultures of cells grown on 5 cm dishes were incubated with 2.4 U/ml Dispase II (Sigma, D4693) in DMEM at 37°C, for 30 min. Cells lifted from the substrate as an intact cell sheet were imaged and then the sheets were submitted to mechanical stress on a shaker at 60 rpm. For measuring the sheet area and counting the sheet fragments, the ImageJ tools (“cell count” and “Polygon selection”) were used.

### Immunofluorescence microscopy

For immunofluorescence, cells were grown for 2 days on glass coverslips or imaging glass-bottom dishes (P35G-1.5; MatTek) and were fixed with 3% formaldehyde (5 min) and then permeabilized with 1% Triton X-100 (15 min), as described previously [31]. The confocal images were taken using a Nikon AXR laser scanning microscope equipped with a Plan Apo 60x×/1.45 objective lens. Immediately before imaging, the dishes were filled with 90% glycerol. The images were then processed using Nikon’s NIS-Elements software. For immunostaining the following antibodies were used: Mouse anti-E-cadherin mAb clones SHE78-7 and HECD1 (Takara, M126 and M106), anti-vinculin (Sigma, V9264), anti-GFP (for Western) and anti-afadin (Santa Cruz Biotechnology, sc-9996 and sc-74433), and anti-p120 (BD Transduction Laboratories, 610134); Chicken anti-GFP (Novus, NB100-1614); Rabbit anti-p120 and anti-α-catenin (Abcam, ab92514 and ab51032, correspondingly), anti-Dsg2 (Proteintech, 21880-I-AP), anti-mCherry (BioVision, 5993-100), anti-PLEKHA5 (Invitrogen, PA5-57463); Guinea pig anti-pkp4 and anti-ARVCF (Progen, GP71 and GP155, correspondingly); and Sheep anti-erbin (R&D systems, AF7866). All secondary antibodies were produced in Donkey (Jackson Immunoresearch Laboratories).

### Live-cell imaging

The live cell imaging experiments were performed essentially as described previously [61] using an X-Cite 120LED Boost High-Power LED Illumination System as a light source. In brief, cells were imaged in L-15 media with 10% FBS by an Eclipse Ti-E microscope (Nikon, Melville, NY) at RT or 37°C controlled with Nikon’s NIS-Elements software. The microscope was equipped with an incubator chamber, a Back illuminated sCMOS Prime-95B camera (Photometrics) and a Plan-Apo TIRF 100x/1.49 lens. No binning mode was used in all live-imaging experiments. At this microscope setting, the pixel size was 110 nm. To monitor the entire lateral membrane from its basal to the apical borders, we obtained stacks of 5 focal planes with 0.5 μm spacing at each time point. All z-stack images were saved in ND2 format. The maximum intensity projection of each frame was created using NIS Element 5.02. For movie analyses, all images were saved as Tiff files and processed using ImageJ software (National Institutes of Health). To track the lateral-sAJ motion, TrackMate plugin of Fiji was used with a setting: DoG detector object diameter of 1 μm, quality threshold of 1, and auto thresholding function. Simple LAP tracker with maximum linking distance of 1 μm, Gap-Closing distance of 1μm and Gap-closing Max frame gap of 1 was used. Tracks were colored based on the Frame origin.

Fluorescence recovery after photobleaching was calculated using Fiji. Initially, several spot-like junctions were selected and precisely bleached with 405 lasers and recovery was documented in every 10 seconds for a total of 500 seconds. For GFP-P120 and GFP-PKP4 recovery was documented in every one second for 100 seconds.

### Platinum Replica Electron Microscopy (PREM)

Sample preparation for immunogold PREM was performed as described previously [63, 64]. In brief, cells grown on glass coverslips were sequentially extracted for 5 min with cytoskeleton preservation buffer (CPB, 100 mM PIPES, pH 6.9; 1 mM EGTA; 1 mM MgCl_2_) supplemented with 1% Triton X-100, then washed with CPB for 1 min, and incubated for 10 min with the anti-mCH antibody in CBP. Then the cells were washed for 5 min in CBP and incubated for additional 10 min with Biotin-SP-VHH fragment Alpaca anti-rabbit antibody (Jackson ImmunoResearch Laboratories, 611-064-215). The staining solutions were supplemented with 1% of Blocking Solution for Gold conjugates (Aurion). After another washing, the cells were fixed with 0.2% glutaraldehyde (for 10 min), quenched with 2 mg/ml NaBH4 in PBS (or 10 min) and stained with 10 nm gold-conjugated Streptavidin (Abcam, ab270041). After final washing (5 min) cells were postfixed with 2% glutaraldehyde in 0.1 M Na-cacodylate, pH 7.3. Fixed cells were treated with 0.1% tannic acid and 0.2% uranyl acetate, critical-point dried, coated with platinum and carbon, and transferred onto EM grids for observation. A parallel coverslip (stained as indicated above together with the control one without the anti-mCH antibody) were fixed in 3% formaldehyde and stained with mouse anti-Streptavidin antibody to verify staining specificity. EM samples were imaged using a FEI Tecnai Spirit G2 transmission electron microscope (FEI Company, Hillsboro, OR) operated at 80 kV. Images were captured by Eagle 4k HR 200kV CCD camera and presented in inverted contrast.

### Data processing

Chemiluminescence was detected via the Azure C300 Chemiluminescent Imager (Azure Biosystems, Dublin, CA) and band intensities were analyzed using ImageJ software (rsb.info.nih.gov/ij/). All other images were processed and analyzed using Nikon’s NIS-Elements ver. 5.02. For line scan analysis, the Element’s in-built line profile function was used to draw a 1pix wide line across the junctions. For peak intensity measurement, the highest intensity on the Y-axis was recorded. A minimum of 15 independent junctions were scanned from five different images. The ratio of the heterochromatic E-cadherin/specific protein-positive pixels versus the total number of the E-cadherin-positive pixels, channels were split and treated with median filter of 1 pixel using Fiji. Both channels were auto-thresholded using Otsu algorithm. Thresholds were selected and added to the ROI manager. ‘AND’ function was used to include the overlapping area from both channels. Percentage of overlap was calculated from the resulting Analysis. For Pearson’s correlation quantification, the images were processed with limited background reduction and denoising function of NIS-Element 5.02 and then Element’s inbuilt colocalization tool was used to obtains the scatterplots of red and green pixel intensities and to evaluate the Pearson’s correlation of the of the entire confocal images. Representative 5 images were analyzed for each staining. The charts and error bars were plotted using GraphPad Prism version 10.2.0. Statistical significance was analyzed using student’s two-tailed t test for two groups. A p value that was less than 0.05 was considered statistically significant.

## Acknowledgments

We thank Drs. T. Svitkina (University of Pennsylvania), for valuable comments and suggestions. Sequencing, flow cytometry and confocal microscopy were performed at the Northwestern University Genetic, Flow Cytometry, and Advanced Microscopy Centers. The authors declare no competing financial interests. The work was supported by National Institute of Health Grant AR070166 and GM148571 (to S.M.T.).

## Figure Legends

**Figure S1.**
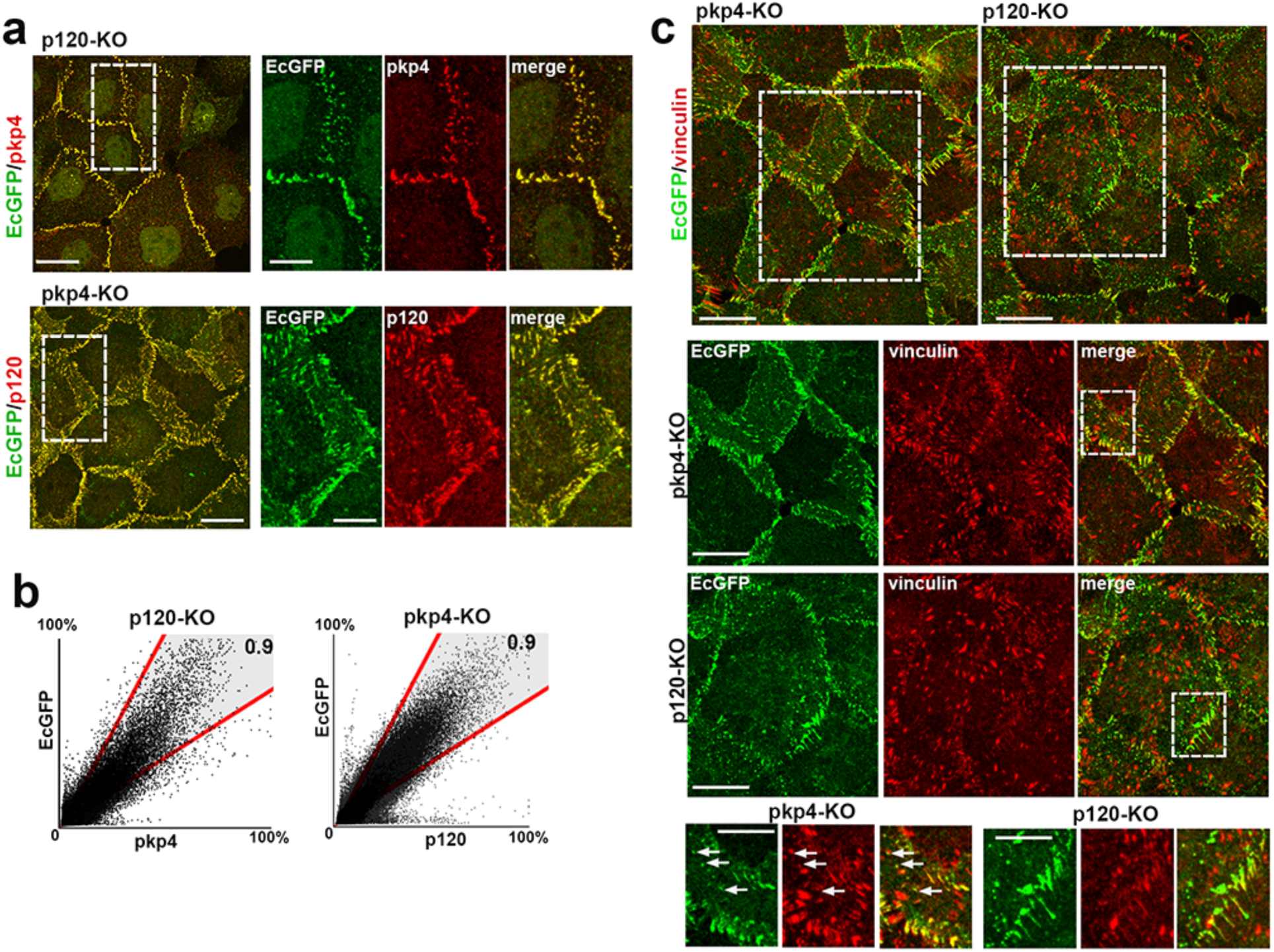
Phenotype of EcGFP-A431 cells deficient for pkp4 and p120. (**a**) Projections of all x-y optical slices of the p120-KP and pkp4-KO cells stained for EcGFP (GFP, green) and pkp4 or p120 (both red) correspondingly. Only merged images are shown for low magnifications. Bar, 20 μm. The zoomed areas (marked by dashed boxes) are shown in both colors at right. Bar, 10 μm. (**b**) The scatterplots of red and green relative pixel intensities of the images shown in a. The r values are at the upper right. Note that the cells show nearly perfect positive relationship between the fluorophores. (**c**) Projections of all x-y optical slices of the KO cells stained for EcGFP (GFP, green) and vinculin (red). Only merged images are shown for low magnifications. Bar, 20 μm. The zoomed areas (marked by dashed boxes) are shown in both colors at the bottom. Bar, 15 μm. Its dashed box area is further zoomed at the bottom row. Bar, 8 μm. The arrows point at vinculin-positive lateral AJs in pkp4-KO cells.

**Figure S2.**
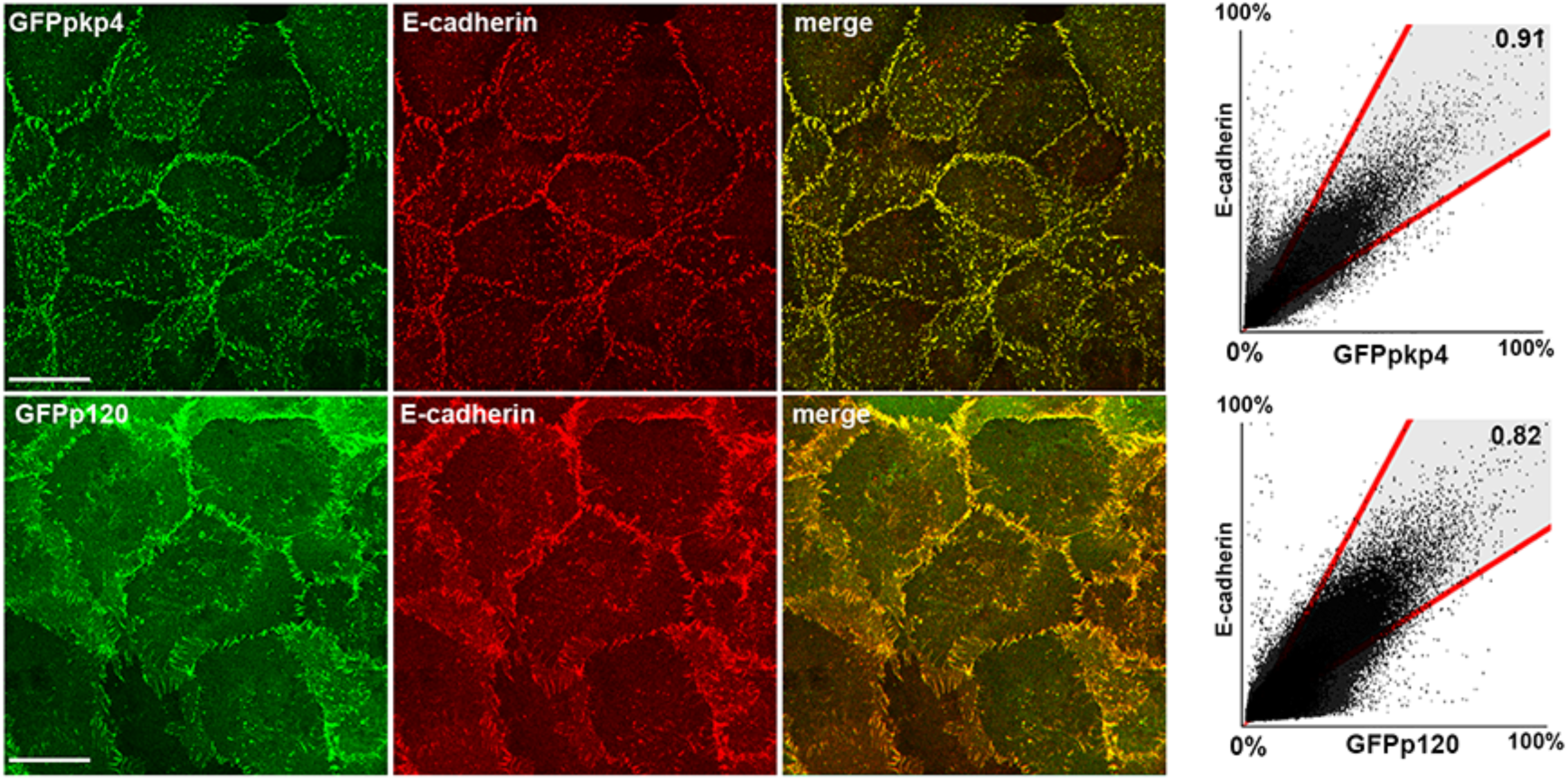
GFPpkp4 and GFPp120 co-localize with E-cadherin. (**a**) Projections of all x-y optical slices of GFPpkp4 and GFPp120 cells deficient for the endogenous p120, pkp4, and ARVCF stained for GFP (green) and E-cadherin (red). Bar, 20 μm. The scatterplots of red and green relative pixel intensities of these images are at the right panel. The r values are at the upper right. Note that the green and red fluorophores show nearly perfect positive relationship in both cells.

**Figure S3.**
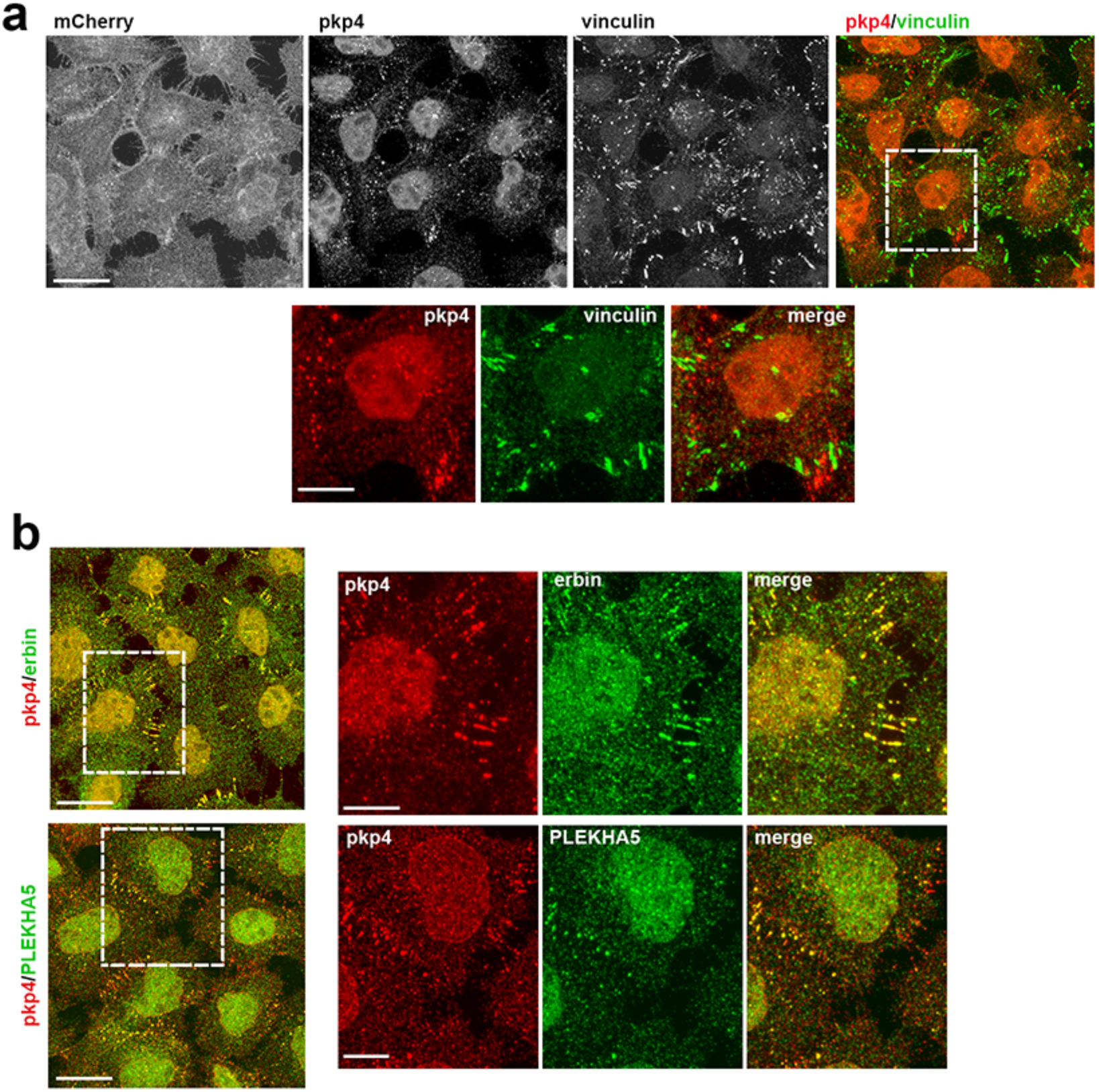
GFPpkp4-E-cadherin clusters display phenotype of the lateral-sAJs. (**a**) Confocal fluorescence microscopy (projections of all x-y optical slices) of αCat-KO cells expressing αCatCH-Δ259 triple stained for mCherry, pkp4, and vinculin. Bar, 20 μm. More detailed subcellular distribution of pkp4 (shown in red) and vinculin (green) of the white dashed boxes area is presented at the bottom. Bar, 10 μm. (**b**) Projections of all x-y optical slices of the same cells double stained for pkp4 (colorized in red) and erbin or PLEKHA5. Only merged images are shown for low magnifications. Bar, 20 μm. The zoomed areas (marked by white dashed boxes) are shown in both colors on the right. Bar, 10 μm.

**Figure S4.**
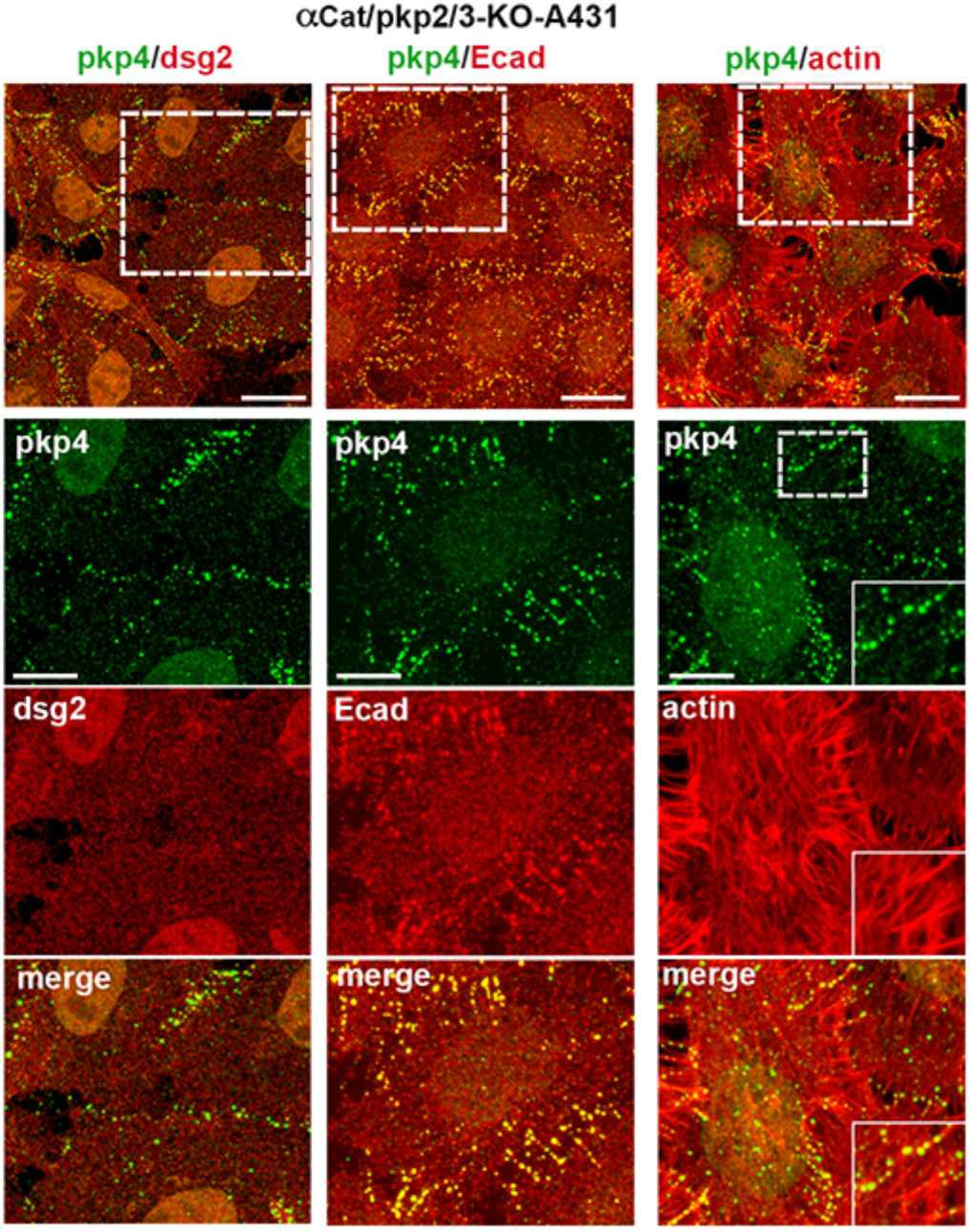
Desmosomes are not involved in pkp4-facilitated E-cadherin clustering. Projections of all x-y optical slices of αCat/pkp2/3-KO cells stained for pkp4 (green) and co-stained for the DSM marker, dsg2, or E-cadherin, or F-actin (all red). Only merged images are shown for low magnifications. Bar, 20 μm. The zoomed areas (marked by white dashed boxes) are shown in both colors at the bottom. Bar, 10 μm. The boxed region within the enlarged image stained for F-actin is further zoomed in the insets. Note that the cells assemble pkp4 clusters despite a complete absence of DSMs. Note also that absolute majority of pkp4 clusters are associated with the actin-rich structures.

**Video S1**. **Dynamics of the AJs at the selected cell-cell contact of the control EcGFP-expressing A431 cells.** The most representative cell-cell contact from 6 independent movies was selected. The images were acquired at 30 sec intervals. See Fig. 3 for details.

**Video S2**. **Dynamics of the AJs at the selected cell-cell contact of the control EcGFP-expressing pkp4-KO cells.** The most representative cell-cell contact from 6 independent movies was selected. The images were acquired at 30 sec intervals. See Fig. 4d for details.

**Video S3**. **Dynamics of the AJs at the selected cell-cell contact of the control EcGFP-expressing p120-KO cells.** The most representative cell-cell contact from 6 independent movies was selected. The images were acquired at 30 sec intervals. See Fig. 4e for details.

**Video S4. Dynamics of the AJs at the selected cell-cell contact of the GFPpkp4-expressing δCat-KO cells.** The most representative cell-cell contact from 8 independent movies was selected. The images were acquired at 30 sec intervals. See Fig. 5e for details.

**Video S5. Dynamics of the AJs at the selected cell-cell contact of the GFPp120-expressing δCat-KO cells.** The most representative cell-cell contact from 8 independent movies was selected. The images were acquired at 30 sec intervals. See Fig. 5f for details.

